# Somatostatin Neurons in the Bed Nucleus of the Stria Terminalis Play a Sex-Dependent Role in Binge Drinking

**DOI:** 10.1101/2022.02.23.481479

**Authors:** Malini Suresh Nair, Nigel C. Dao, Daniela Lopez Melean, Keith R. Griffith, W. David Starnes, J. Brody Moyer, Avery R. Sicher, Dakota F. Brockway, Kathleen D. Meeks, Nicole A Crowley

## Abstract

Alcohol use disorder (AUD) is characterized by alcohol use coupled with chronic relapse and involves brain regions including the bed nucleus of the stria terminalis (BNST). Here, we explore whether a subpopulation of BNST neurons, somatostatin (SST) expressing GABAergic neurons, play a role in an animal model of binge-like alcohol consumption, the Drinking in the Dark (DID) model. Chemogenetic activation of BNST SST neurons reduced binge alcohol consumption in female but not male SST-Cre mice, while inhibition of these neurons in the same mice had no effect. In addition, chemogenetic activation of these neurons did not cause apparent changes in models of anxiety-like behavior in either sex. Basal SST cell counts and intrinsic excitability of SST neurons were compared to attempt to understand sex differences in DREADD-induced changes in drinking, and while males had a greater number of BNST SST neurons, this effect went away when normalizing for total BNST volume. Together, these results suggest SST neurons in the BNST should be further explored as a potential neuronal subtype modulated by AUD, and for their therapeutic potential.

**HIGHLIGHTS:**

- Chemogenetic activation of BNST SST neurons reduces binge drinking in female but not male mice
- Chemogenetic activation of these neurons has no effect on anxiety-like behavior in either sex
- Electrophysiology revealed no clear sex differences in intrinsic excitability BNST SST neurons between males and females
- Imaging revealed males had greater overall BNST SST cell numbers than females, but this effect could be explained by normalizing for total BNST volume

## 1. INTRODUCTION

Recent estimates found that alcohol binge drinking, defined as rapid, excessive alcohol consumption, was projected to cost the U.S. approximately 191.1 billion dollars per year - amounting to 70% of the total societal cost of alcohol drinking (Sacks et al., 2015). Binge drinking behaviors create a cascade of large and negative economic and health effects for those who partake in it, manifesting in both social (lost employment, relationships) and personal (cancer, other illnesses) domains (Alcohol Use Disorder, 2018). Repeated binge drinking behavior can lead to alcohol dependence that increases the risk of developing major depressive disorder (MDD) by four-fold (Hasin & Grant, 2002) as well as an anxiety disorder (Grant et al., 2004) - setting the stage for cascading cycles of relapse to alcohol and greater mental health issues. Finding neuronal targets to reduce binge drinking will play a critical role in ameliorating both AUD and related neuropsychiatric conditions.

The clinical and animal literature has suggested somatostatin (SST) as a promising target for multiple neuropsychiatric disorders. SST expressing neurons are found throughout the brain, including in the amygdalar regions. The bed nucleus of the stria terminalis (BNST) is a region of the extended amygdala that is highly interconnected with other regions such as the medial prefrontal cortex (mPFC, Lebow, M. A., & Chen, A., 2016; Crowley et al., 2016) and the ventral tegmental area (VTA; Jennings et al., 2013). The BNST, and the amygdala more broadly, are implicated in the withdrawal and negative affect stage of drug addiction (Koob & Volkow, 2010; Kash et al., 2015; Pleil et al., 2015). Our lab and others have shown alcohol influences SST neurons in regions of the mPFC (Joffe, Winder, & Conn, 2020; Dao et al., 2021; Li et al., 2021) - though little has been done to characterize BNST SST neurons and alcohol. Previous work by our lab showed that BNST SST neurons display decreased intrinsic excitability following a forced abstinence model (Dao et al. 2020). This work, modeled off known interactions between the BNST and intermittent access to alcohol (Holleran et al., 2016), characterized the connection between intermittent access to alcohol, withdrawal, and an acute stressor (forced swim), but importantly, did not explore alcohol alone. Others have similarly comprehensively explored SST neurons in the central amygdala and BNST pathway in anxiety-like behavior (Ahrens et al., 2018), but did not explore substance use.

The following experiments aimed to elucidate the role of SST BNST neurons in binge alcohol consumption behavior and anxiety-like behavior in order to establish preclinical evidence for their efficacy as a therapeutic target.

## 2. MATERIALS AND METHODS

### 2.1 Mice

All experiments were approved by the Pennsylvania State University Institutional Animal Care and Use Committee. Male and female hemizygous SST-IRES-Cre mice (stock #007909, The Jackson Laboratory) and Ai9 reporter mice (stock #007909, The Jackson Laboratory) on a C57BL/6J background (stock #000664, The Jackson Laboratory) were bred in house under normal conditions. Adult mice (over 6 weeks of age) were used for all experiments, and mice were single-housed on a 12 hr reverse light cycle for (lights off at 7:00 am) for a minimum of one week prior to any behavioral manipulations. All mice had *ad libitum* access to food and water, with the exception that water was removed during the DID procedure.

### 2.2 Stereotaxic surgery for viral injections

Starting at 8 weeks of age, mice underwent intra-BNST (from Bregma, AP: +0.3 mm; ML: +/- 1 mm, DV: −4.30 mm) injections of viral constructs. Mice were anesthetized with isoflurane (5% induction, 1-2% maintenance) and mounted on the stereotaxic frame (Stoelting, Wood Dale, IL). Following craniotomy, 0.3-0.4 uL of virus was injected in each hemisphere of the BNST at a rate of 0.1 uL/min via a 1 uL Hamilton Neuros Syringe. Post-injection, the syringe was left in place for 5-6 min before being removed slowly to limit efflux of virus from the injection site. For postoperative pain management, ketoprofen (5mg/kg) and bupivacaine (4mg/kg) were applied intraperitoneally and topically, respectively. Following surgeries, mice were monitored and allowed to recover for one week. To allow for sufficient viral expression, experimental manipulations (Designer Receptors Exclusively Activated by Designer Drugs, DREADDs; ligands administration or electrophysiology) were performed at least 3 weeks out from viral injections.

### 2.3 Viruses

Viral vectors AAV8-hsyn-DIO-hM3D-(Gq)mCherry (Armbruster et al., 2007; Alexander et al., 2009), AAV8-hsyn-dF-MA-KORD-IRES-mCirine (Vardy et al. 2016), and AAV5-syn-DIO-mCherry were obtained from Addgene (Watertown, MA). For alcohol DID and sucrose DID experiments, the hM3D and KOR DREADD viruses were cocktailed to drive both excitation and inhibition of SST neurons in the same animals. For anxiety-like behaviors, only the excitatory DREADD was tested due to limitations around repeated testing in anxiety-like behaviors.

### 2.4 Alcohol Drinking in the Dark (DID)

DID was conducted as it is described in previous publications (Rhodes et al., 2005; Crowley et al., 2019a; Crowley et al., 2019b). Mice received 20% (v/v) ethanol (EtOH; Koptec, Decon Labs, King of Prussia, PA) for 2 hr in drinking water, 3 hr into the dark cycle (10 am) on three sequential days. Mice received 20% ethanol for 4 hr on the fourth day (also known as the binge day). After the binge day, mice underwent three days of abstinence before repeating the DID cycle (the number of cycles per experiment is described below). Mice underwent a week of baseline DID before surgeries.

### 2.5 Sucrose Drinking in the Dark (DID)

In both cohorts, four days after the last EtOH binge session, mice underwent sucrose DID conducted as it is described previously (Rinker et al., 2017). Mice received 10% sucrose in water (w/v) 3 hr into the dark cycle on three sequential days. Mice received 10% sucrose for 4 hr on the fourth day (i.e., the alcohol DID and sucrose DID protocols were run identically with the exception of substance consumed). Each cohort underwent three cycles of sucrose DID.

### 2.6 Anxiety-like Behavior

Mice were tested for anxiety-like behavior using standardized procedures (Crowley et al 2016; Al-Hasani et al 2015), using the elevated plus maze (EPM) and the open field test (OFT). Behaviors were run three weeks following stereotaxic injection of viral constructs. Prior to undergoing each test, mice were allowed to rest for at least 1 hr in the testing room. Both tests were done 3 hr into the dark cycle and under red light (6 lux). Mice received injections of CNO (3 mg/kg; i.p) 30 mins before undergoing each behavioral test (spaced 3 days apart).

#### Elevated Plus Maze

For the EPM, each mouse was placed in the center of the square maze (30 x 5 x 40 cm), facing a closed arm (20 cm arm wall height, transparent Plexiglass and gray floor). Each mouse was allowed to explore the EPM maze for 5 min. All sessions were recorded with EthoVision XT video tracking system (Noldus, Leesburg, VA, United States), which automatically analyzed the total time spent in the open arms and the number of entries into the open arms.

#### Open Field Test

For the OFT, each mouse was placed in a corner of a black Plexiglass arena (50 x 50 x 20 cm) and left to explore for 20 min. Once again, sessions were recorded with EthoVision XT which analyzed the total time spent in the center zone, as well as the number of entries to the center zone. The center zone was defined as a 12.5 x 12.5 cm area at the center of the OFT arena.

### 2.7 Drug Administration

30 mg/ml clozapine-*n*-oxide (CNO; #HB1807, HelloBio, Princeton, NJ) was dissolved in pure DMSO (Fisher Scientific, Waltham, MA) and diluted to 0.3 mg/ml in 0.9% saline. 10 mg/ml of salvinorin-B (SalB; HB4887, HelloBio, Princeton, NJ) was dissolved in pure DMSO. For DID experiments, mice were habituated to handling and injections and received 0.1 ml/g bodyweight of sterile saline on cycle 2. Mice received vehicle injections of either 1% DMSO in saline (10 ml/kg; intraperitoneal, i.p.) or pure DMSO (1 ml/kg; subcutaneously, s.c.) on day 2 of DID cycles 3 and 4. All mice receiving CNO and SalB, received these control injections of 1% DMSO and pure DMSO, respectively, as these corresponded to the solvent into which each drug was dissolved to achieve the desired concentrations at that particular week. Injections of CNO and SalB were counterbalanced across mice to account for any order effects that might occur (i.e., mice that received CNO (3 mg/kg; i.p.) on cycle 3, received SalB (10 mg/kg; s.c.) on cycle 4 and vice-versa). For sucrose DID, mice received vehicle injections on day 4 of cycle 1, and CNO or SalB injections on cycles 2 and 3. For anxiety-like behavior, mice were habituated to handling and injections, as described previously, every day, 3 days leading up to behavior testing. Mice underwent both EPM and OFT (3 days apart) and received the same dose of CNO as previously described.

### 2.8 Electrophysiology

Confirmation of DREADD expression and function was validated using electrophysiology in a subset of mice, as described previously. Experiments were conducted post-DID exposure. Mice were deeply anesthetized via inhaled isoflurane and rapidly decapitated. Brains were quickly removed and processed according to the NMDG protective recovery method (Ting et al., 2018). Briefly, brains were immediately placed in ice-cold oxygenated N-methyl-D-glucamine (NMDG)-HEPES aCSF containing the following, in mM: 92 NMDG, 2.5 KCl, 1.25 NaH2PO4, 30 NaHCO3, 20 HEPES, 25 glucose, 2 thiourea, 5 Na-ascorbate, 3 Na-pyruvate, 0.5 CaCl2·2H2O, and 10 MgSO4·7H2O (pH to 7.3-7.4). The BNST was identified according to the Allen Mouse Brain Atlas. 300 μM coronal slices containing the BNST were prepared on a Compresstome vibrating microtome (VF-310-0Z, Precisionary Instruments LLC, Greenville, NC, United States), and transferred to heated (31°C) NMDG-HEPES aCSF for a maximum of 10 min. Slices were then transferred to heated (31°C) oxygenated normal aCSF (in mM: 124 NaCl, 4.4 KCl, 2 CaCl2, 1.2 MgSO4, 1 NaH2PO4, 10.0 glucose, and 26.0 NaHCO3, pH 7.4, mOsm 300-310), where they were allowed to rest for at least 1 h before use. Finally, slices were moved to a submerged recording chamber (Warner Instruments, Hamden, CT) where they were continuously perfused with the recording aCSF (2 ml per min flow rate, 31°C). Recording electrodes (3-6 MΩ) were pulled from thin-walled borosilicate glass capillaries with a Narishige P-100 Puller (Amityville, NY, United States). All experiments used a potassium-gluconate-based (KGluc) intracellular recording solution, containing the following (in mM): 135 K-Gluc, 5 NaCl, 2 MgCl2, 10 HEPES, 0.6 EGTA, 4 Na2ATP, and 0.4 Na2GTP (287-290 mOsm, pH 7.35).

CNO (10 μM) or SalB (100 nM) was added to the recording aCSF for 10 min following a 10 min stabilization of resting membrane potential (RMP), and the recording continued for a 15 min washout. Tetrodotoxin (500 nM) was added to the recording aCSF to block action potentials for consistent measurement of RMP. Average RMPs were normalized to a 5 min baseline.

In a separate behavior-naive cohort of mice, intrinsic excitability was compared in Ai9-expressing SST cells in the BNST of male and female mice. SST-expressing neurons were identified in SST-IRES-Cre::Ai9 mice via presence of tdTomato fluorescence under a 40× immersed objective with 565 nm LED excitation. Measurements of intrinsic excitability included resting membrane potential (RMP), rheobase (the minimum amount of current needed to elicit an action potential during a current ramp protocol), action potential threshold (the membrane potential at which the first action potential fired), and the number of action potentials fired during a voltage-current plot protocol (V-I plot) with increasing steps of depolarizing currents (0-200 pA, 10 pA per step). Hyperpolarizing currents (not shown) were included as a control. Experiments were performed at both RMP and at the standard holding potential of −70 mV.

### 2.9 Histology and Imaging

Mice were deeply anesthetized with Avertin (250 mg/kg) and transcardially perfused, first with ice-cold phosphate (PBS) followed by 4% (w/v) paraformaldehyde (PFA). Brains were post-fixed in PFA overnight and sectioned at 40-60 microns using a Compresstome vibrating microtome (VF-300-0Z, Precisionary Instruments LLC, Greenville, NC, United States) or a Leica vibratome (VTS 1200, Leica). BNST-containing sections were mounted on SuperFrost glass slides air-dried, and coverslipped with ImmunoMount (Thermo Fisher Scientific, Waltham, MA, United States) mounting media. Viral injections were assessed under mCitrine or mCherry fluorescence filters on an Olympus BX63 upright microscope (Center Valley, PA), and the brightest fluorescence point was chosen as the center of the injection. While control mice with unilateral expression were included in this data set, experimental mice with unilateral expression or missed injections were excluded from data analysis.

### 2.10 SST Cell Counts

SST+ cell counts were quantified by researchers blind to mouse sex using ImageJ (National Institutes of Health, Bethesda, MD, United States). The region of interest (ROI), either the dorsal BNST (dBNST) or the ventral BNST (vBNST, not shown), was delineated and SST+ were automatically quantified under matched criteria for size, circularity and intensity consistently with our previously published work (Dao et al. 2020). Threshold to highlight all of the SST+ cells counted was set between 3-4%. Each ROI's total SST cell count was divided by the ROI area to give a total SST+ density value (Smith et al., 2019).

### 2.11 Statistical Analysis

Experimenters were blinded to group (DID or water), and viral construct (DREADD or control vector) whenever possible. In EtOH DID and sucrose DID experiments, data were analyzed by mixed-model ANOVAs with virus or sex as between-subject factors and drug treatments as within-subject factor. Despite the differences in vehicles for preparation of CNO and SalB (1% DMSO in saline vs. pure DMSO) and in administration routes (intraperitoneal vs. subcutaneous), there was no significant differences in week 2 (vehicle injection week) alcohol consumption levels (Student's *t*(19) = 0.385, *p* = 0.704) and so vehicle data were collapsed. Bonferroni's post-hoc tests were used for pairwise between-group comparisons, whereas Dunnett's post-hoc tests were used for within-group comparisons between vehicle administrations and CNO or SalB administrations. EPM and OFT data were analyzed by ordinary two-way ANOVAs. Effects of CNO and SalB on SST neurons' RMP was analyzed using paired *t*-test (5-min washout vs. 5-min baseline) and Student's *t*-test. Statistical analysis and graph construction was performed in Graphpad Prism 7.0 and BioRender. Data are presented as Means and Standard Error of the Mean (SEM). All group *n* per sex, means and SEMs are presented in **Table 1**.

**Table 1.**
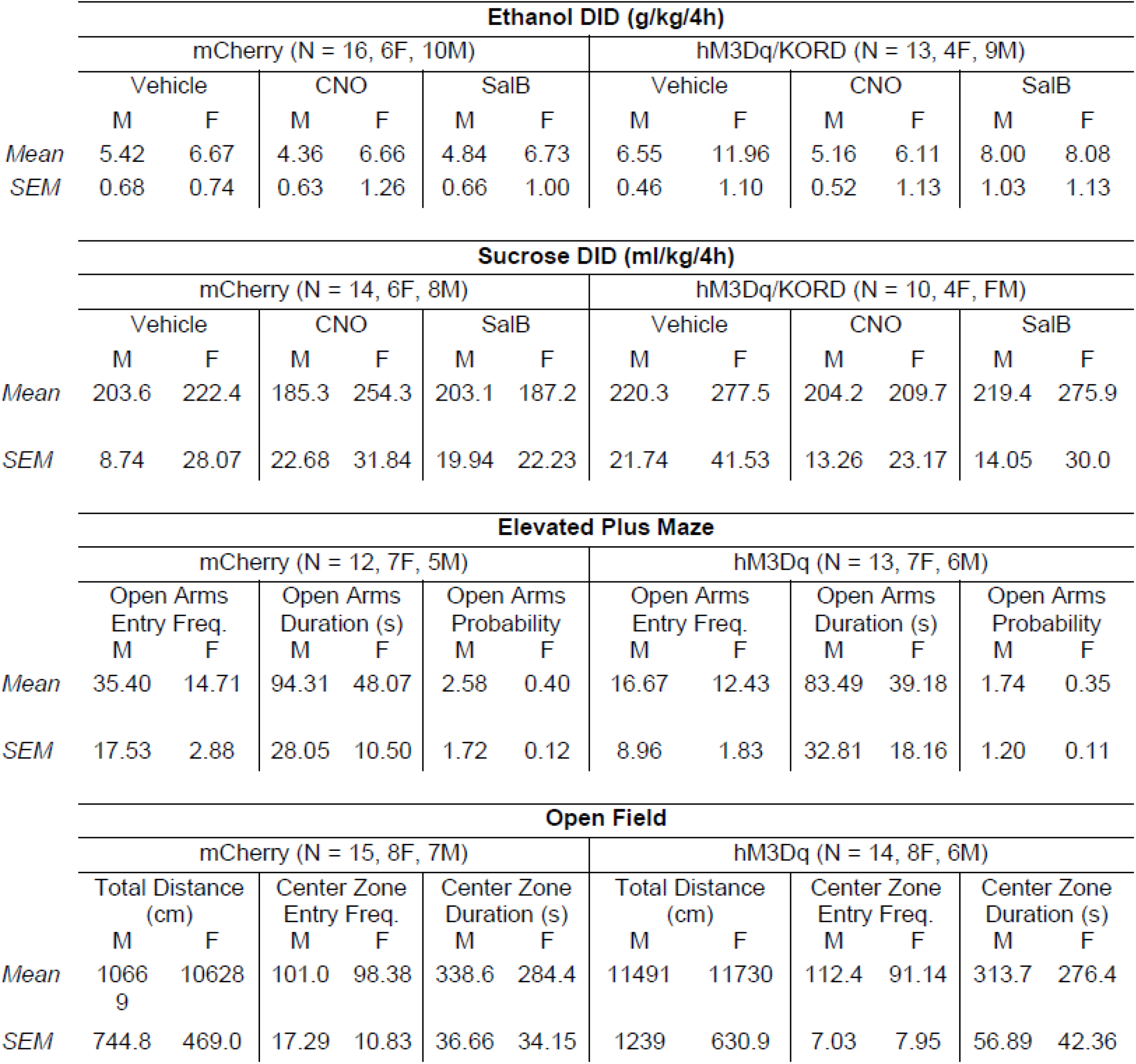
Means and standard errors of the means (SEM) of behavior experiments.

## 3. RESULTS

### 3.1 Activation of BNST SST neurons reduced binge drinking in female, but not male, mice

Adult male and female SST-IRES-Cre mice underwent chemogenetic manipulations of SST neurons to assess their role in binge drinking (**Figure 1A** for timeline, **Figure 1B** for representative injections). In DREADD expressing mice, CNO activation of SST neurons reduced binge drinking, while SalB inactivation of SST neurons had no effect on alcohol consumption levels (n = 16 mice for mCherry controls (6F, 10M) and 13 mice for hM3Dq/KORD (4F, 9M), F_virus_(1, 27) = 4.601, *p* = 0.041, F_treatment_(2, 54) = 7.547, *p* = 0.001, F_virus x treatment_(2, 54) = 4.201, *p* = 0.020, CNO vs. Vehicle: *p* < 0.001, SalB vs. Vehicle: *p* > 0.1), **Figure 1C**). Neither CNO nor SalB (*p* > 0.1) administration had any effect on binge drinking in mice injected with the control virus mCherry, ruling out gross off-target effects. SalB-injected hM3Dq/KORD mice also drank more than SalB-injected mCherry controls (*p* = 0.017). In hM3Dq/KORD mice only, the effect of SST neurons activation and inhibition were evident only in females (F_sex_(1, 11) = 5.756, *p* = 0.035, F_treatment_(2, 22) = 9.726, *p* < 0.001, F _sex x treatment_(2, 22) = 5.865, *p* = 0.009, **Figure 1D**). Again, CNO administration reduced alcohol consumption levels in females (CNO vs. Vehicle: *p* < 0.001), but there was also a small but significant reduction in alcohol consumption levels following SalB administration (SalB vs. Vehicle: *p* = 0.020), reminiscent of the effects we previously reported in prelimbic SST neurons (Dao et al., 2021). In contrast, neither CNO nor SalB was able to reduce alcohol consumption in males (*p* > 0.1).

**Figure 1:**
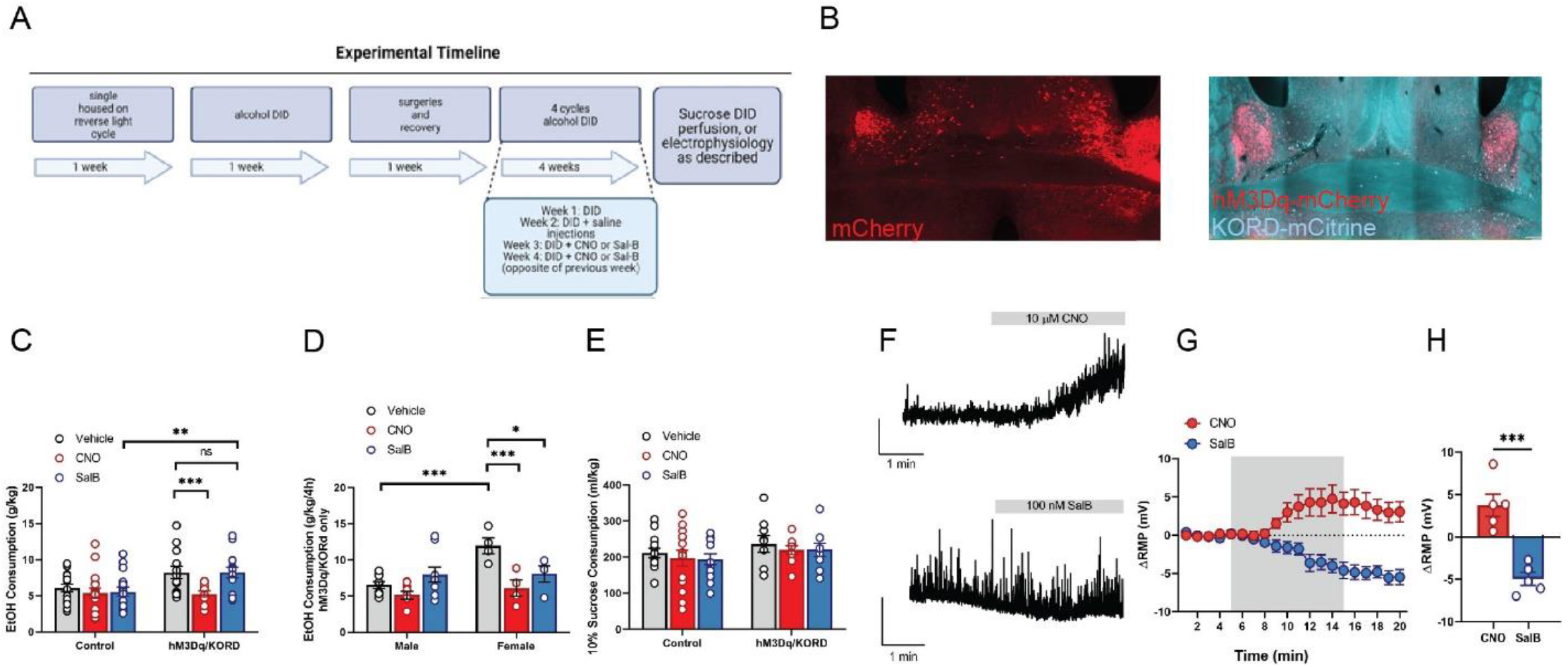
Chemogenetic activation of BNST SST neurons reduced binge drinking. **(A)** Timeline of experimental manipulations. **(B)** Representative images of mCherry control injections (*left*) and KOR-DREADD/hM3D multiplexed injections (*right*). Multiplexed DREADD manipulations had high fidelity of coexpression. **(C)** Chemogenetic activation of SST neurons via activation of Gq-coupled DREADD reduced binge drinking, whereas neither CNO nor SalB had any effects in control (non-DREADD expressing) mice. **(D)** In hM3Dq/KORD-expressing mice, the effect of CNO administration were only observed in females but not males. There was also a small but significant reduction of alcohol binge drinking following SalB administration in females. **(E)** There was no effect of activation of either DREADD on sucrose consumption in a sucrose DID paradigm. For **C-D**, filled circles represent female data, and blank circles represent male data. **(F)** Representative RMP traces during bath application of CNO (10 μM, *top*) and SalB (100 nM, *bottom).* **(H-G)** As expected, CNO depolarized and SalB hyperpolarized the RMP of DREADD-expressing SST neurons. The average change in RMP was calculated during washout (min 16-20).

We then explored whether similar effects were seen following a sucrose DID model (**Figure 1E**). Manipulation of SST neurons did not alter sucrose consumption in either sex (F_virus_(1,21) = 1.9, *p* = 0.183; F_treatment_(2, 42) = 0.639, *p* = 0.533; F_virus x treatment_(2, 42) = 0.0; *p* = 0.991). Electrophysiology was used to confirm DREADD functionality in a sub-cohort of mice (Figure **Figure 1 F-H**). As excepted, bath application of CNO depolarized the RMP of DREADD-expressing SST neurons (wash-out vs. baseline: paired *t*(4) = 2.790, *p* = 0.049), while bath application of SalB hyperpolarized the RMP of DREADD-expressing SST neurons (washout vs. baseline: paired *t*(4) = 6.467, *p* = 0.003; CNO vs. SalB: unpaired *t*(8) = 5.622, *p* < 0.001).

### 3.2 Activation of BNST SST neurons had no effect in anxiety-like behavioral assays

We first began by assessing Gq-coupled DREADD signaling as chemogenetic activation showed effects on binge drinking behavior (**Figure 2A** for timeline, **Figure 2B** for representative injections). Animals underwent both the EPM and OF in a counter-balanced order. No significant effects were seen in any measurements of the EPM test (**Figure 2C** representative heat maps; n = 12 mCherry mice (7M, 5F), n = 13 hM3Dq mice (7F, 6M), 3 mice were removed from the control group and 2 from the DREADD group for jumping off the maze or unilateral viral expression). There were no significant differences between virus groups and sexes seen in open arm entrance frequency (F_sex_(1, 21) = 2.210, *p* = 0.152, F_virus_(1, 21) = 1.571, *p* = 0.224, F _sex x virus_(1, 21) = 0.962, *p* = 0.338, **Figure 2D**), open arm duration (F_sex_(1, 21) = 3.969, *p* = 0.060, F_virus_(1, 21) = 0.188, *p* = 0.669, F_sex x virus_(1, 21) = 0.00, *p* = 0.967, **Figure 2E**), open arm entry probability (F_sex_(1, 21) = 3.947, *p* = 0.060, F_virus_(1, 21) = 0.239, *p* = 0.630, F_sex x virus_(1, 21) = 0.198, *p* = 0.661, **Figure 2F**). Mice that underwent OF testing (n = 15 mCherry (8F, 7M), n = 14 hM3Dq (8F, 6M) showed little effects with SST neuronal activation. There were no differences seen in total distance traveled (F_sex_(1, 25) = 0.017, *p* = 0.897, F_virus_(1, 25) = 1.591, *p* = 0.219, F_sex x virus_(1, 25) = 0.033, *p* = 0.855, **Figure 2H**), indicating that it is unlikely BNST SST neuronal activation produced gross motor changes in any of the assays. There was also no significant effect in OF center zone entrance frequency (F_sex_ (1, 25) = 1.082, *p* = 0.308, F_virus_(1, 25) = 0.033, *p* = 0.857, F_sex x virus_(1, 25) = 0.658, *p* = 0.425, **Figure 1I**) and center zone duration (F_sex_(1, 25) = 1.138, *p* = 0.296, F_virus_(1, 25) = 0.146, *p* = 0.705, F_sex x virus_(1, 25) = 0.039, *p* = 0.844, **Figure 1J**).

**Figure 2:**
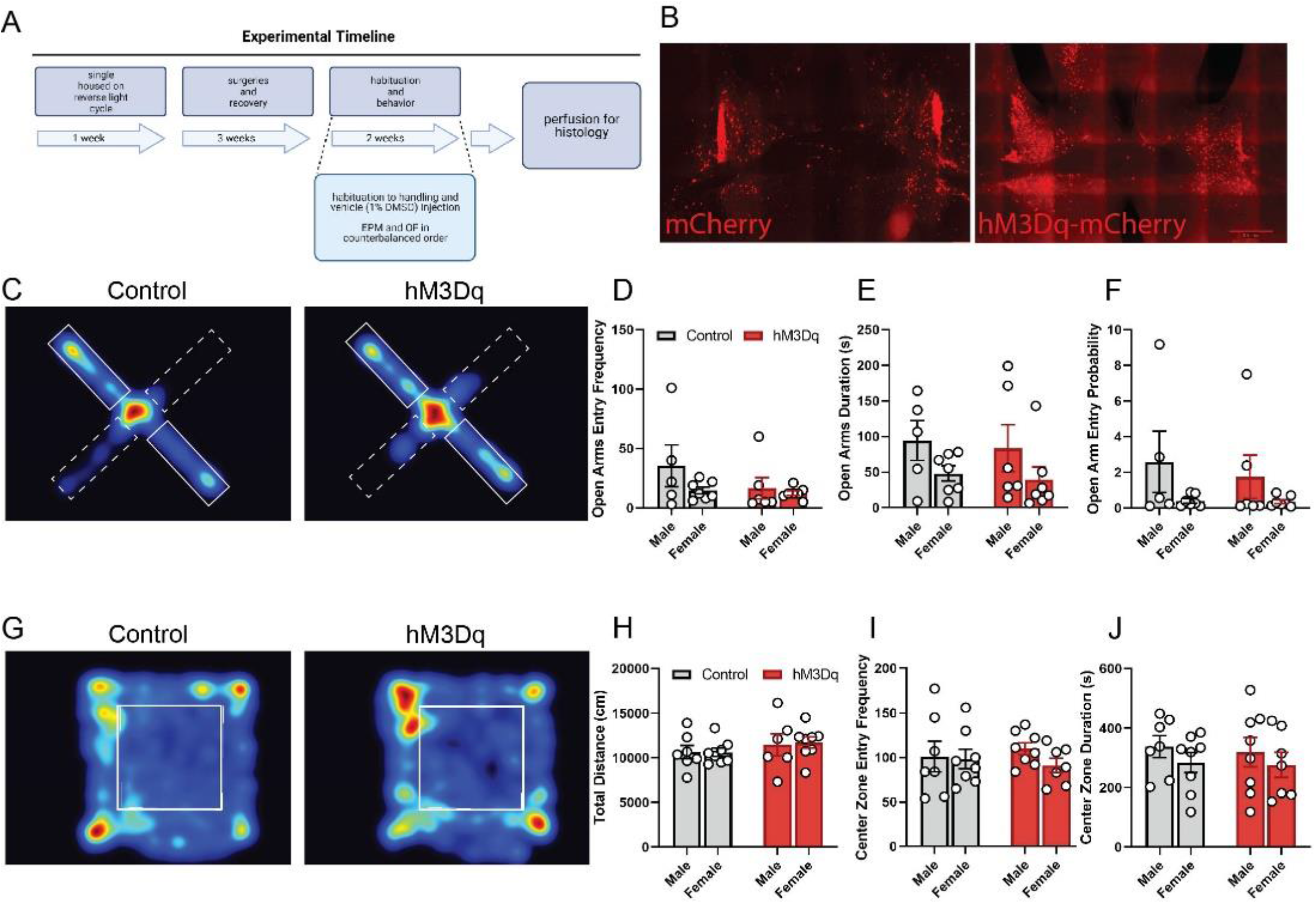
chemogenetic activation of BNST SST neurons showed no substantial effects in the elevated plus maze and open field. (A) Timeline of experimental manipulations. **(B)** representative image of mCherry control (*left*) and hM3D (*right*) injections in the BNST. **(C)** Representative heat maps showing time spent in the EPM open arms (dotted rectangles) and closed arms (full rectangles) in control virus-expressing mice and hM3Dq-expressing mice. **(D)** hM3Dq-induced activation of BNST SST neurons did not change open arms entry frequency, **(E)** open arms duration, **(F)** or open arms entry probability. **(G)** Representative heat maps of time spent in overall regions of the OF. (**H)** hM3Dq-induced activation of BNST SST neurons did not change total distance traveled, numbers of entry into center zone **(I)** or time spent in center zone **(J).**

### 3.3 Comparison of SST intrinsic excitability and total SST cell number in the BNST of males versus females revealed no differences between the sexes

We next sought to understand why chemogenetic activation of BNST SST neurons produced divergent effects on binge drinking in male and female mice. As the BNST and amygdala more broadly is known to be sexually dymorphic, and previous work from our group showed an interaction between alcohol, stress, and SST neurons in the BNST (Dao et al., 2019) we compared intrinsic excitability and total SST cell numbers in the dorsal BNST of male and female SST-Ai9 reporter mice. The intrinsic excitability (representative traces **Figure 3A**) revealed no clear differences between the sexes. BNST SST cells in females and males showed no significant differences in resting membrane potential (*t*(16) = 0.413, *p* = 0.685; **Figure 3B**), rheobase at RMP (*t*(16) = 0.843, *p* = 0.411; **Figure 3C**), action potential threshold at RMP (*t*(16) = 0.011, *p* = 0.991; **Figure 3D**), rheobase at −70 mV (*t*(16) = 1.355, *p* = 0.195: **Figure 3E**), or action potential threshold at −70 mV (*t*(15) 1.017, *p* = 0.325; **Figure 3F**). We also saw no significant differences in the number of action potentials fired per positive current injection at either RMP (2-way ANOVA revealed an expected effect of current injection, F(1.592, 25.47) = 11.92, *p* < 0.001. but no effect of sex F(1,16) = 0.129, *p* = 0.723 and no interaction F(20,320) = 0.126, *p* > 0.999; **Figure 3G**) or −70 mV (again there was an expected significant effect of current injection, F(1.553, 24.85) = 13.53, *p* < 0.001, but not of sex F(1,16) = 1.139, *p* = 0.301 or interaction F(20,320) = 0.438, *p* = 0.984; **Figure 3H**). We next explored whether overall number of SST-expressing cells in the dorsal BNST differed between males and females (representative pictures **Figure 3I**). Though we saw changes in cell count when not normalizing for BNST total size between males and females (effect of sex F(1,8) = 18.94, *p* = 0.002, effect of region F(1,8) = 9.567, *p* = 0.015, interaction F(1,8) = 1.582, *p* = 0.244). **Figure 3J**), no significant differences were seen in cell count per total area when normalized (**Figure 3K**).

**Figure 3:**
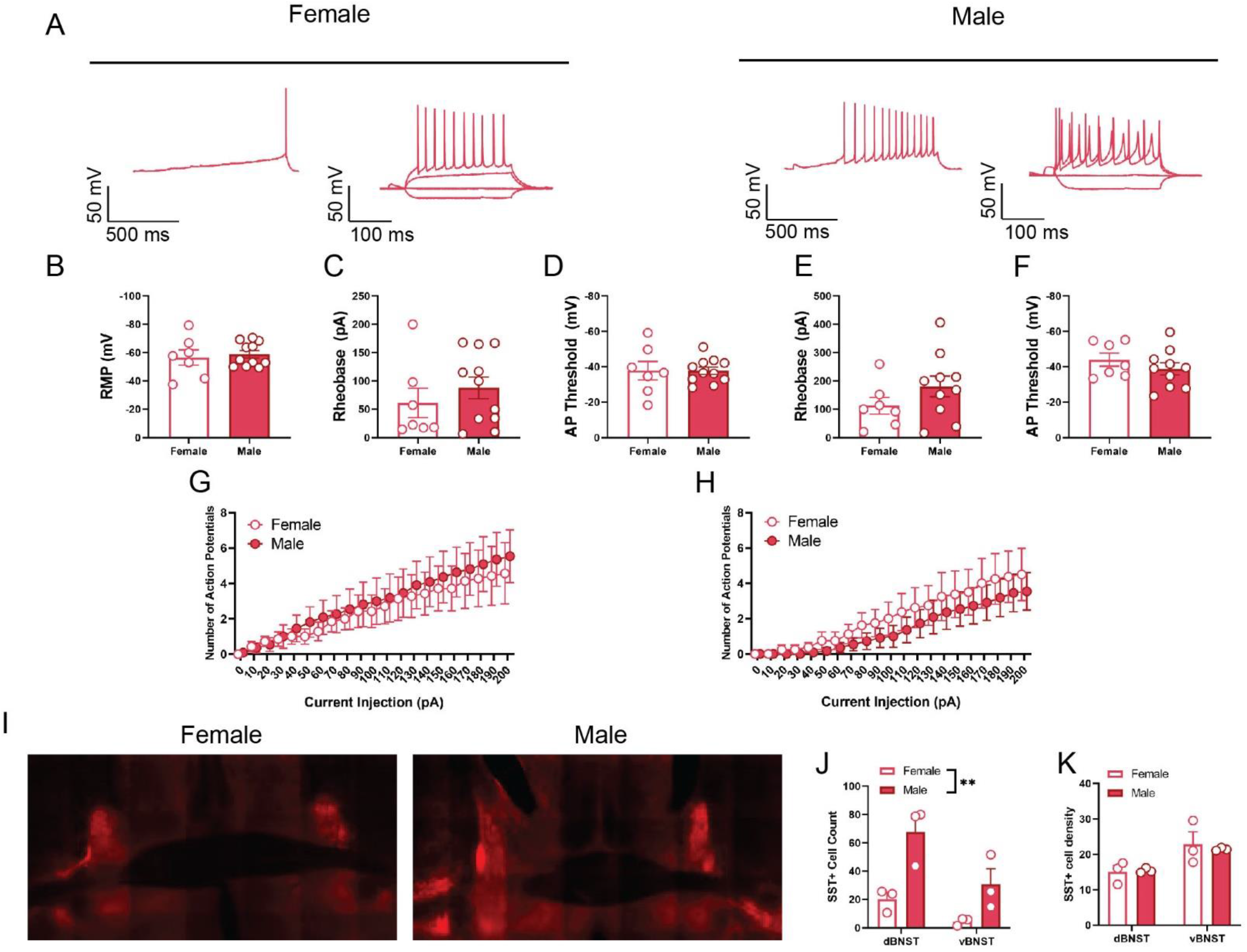
Basal characterization of intrinsic excitability and overall number of SST neurons in the dorsal BNST of male and female reporter mice. **(A)** Representative traces of rheobase and VI plots for males and females at RMP and −70 mV. No significant differences were seen for any of the intrinsic excitability measurements conducted, including comparison of resting membrane potential **(B)**, rheobase at RMP **(C)**, action potential threshold at RMP **(D),** rheobase at −70 mV **(E)**, and action potential threshold at −70 mV **(F)**. We also saw no difference between the sexes in action potentials fired by current injection step at either RMP **(G)** or −70 mV (H). While we saw a greater number of SST neurons in both the dorsal and ventral BNST of male mice as compared to females, SST cell number did not differ when accounting for total area (representative images of Ai9-expressing mice in **I**, quantification in **J-K**).

Together, these experiments provide early evidence that BNST SST neurons may play a role in alcohol consumption behaviors, without interacting with anxiety-like behaviors. This effect did not appear to be due to gross morphological differences or differences in intrinsic excitability between female and male mice.

## 4 DISCUSSION

### 4.1 Overall discoveries and limitations

The goals of the current study were to understand the behavioral effects of manipulating BNST SST neurons. Using a multiplexed chemogenetic approach, we found that activation of this population of neurons reduced binge drinking in female, but not male, mice using the DID protocol. These effects were seen following multiple weeks of baseline drinking (allowing for control injections for DREADD manipulations), however intake levels were relatively stable across weeks (consistent with our previous publications such as Dao et al. 2021 and those published by others (Rinker et al., 2017). Future work should investigate whether chemogenetic manipulation of BNST SST neurons has an effect on alcohol drinking concurrent with the first exposure to alcohol. In addition, though we did not see an effect of KOR-DREADD mediated inhibition of SST neurons on drinking, this does not preclude potential changes with greater inhibition. While we considered an increased dose of SalB (and DMSO) a technical limitation, future work should explore Gi/o coupled DREADDs or optogenetic approaches.

The current work did not explore using the multiplexed DREADD approach due to limitations in repeat testing in the anxiety-like behavioral assays. Interestingly, while we found no effects of chemogenetic manipulation of these neurons on measurements of anxiety-like behaviors, it would be prudent for further research to explore inhibition of these circuits. We also explored a relatively limited range of anxiety-like behavioral assays (open field and elevated plus maze) and a more thorough characterization may reveal a nuanced role of BNST SST neurons on anxiety-like behaviors.

### 4.2 BNST circuitry and alcohol

Both activation and inactivation of BNST neurons (through chemogenetics, optogenetics, or pharmacology) can influence alcohol consumption depending on the sub-population and precise circuitry manipulated. In addition, effects are dependent on the alcohol model used (chronic versus acute, forced versus voluntary, etc), and importantly, work done using different models of ethanol exposure or patterns may not produce consistent effects. For example, broadly silencing the BNST in male DBA/2J mice lead to a reduction in ethanol seeking (Pina et al., 2015). Similarly, silencing VTA-projecting GABAergic neurons with the dorsolateral BNST reduced ethanol consumption (Companion & Thiele, 2018). This publication demonstrates the VTA-targeting neurons are largely corticotrophin-releasing factor (CRF) expressing (87%) and aligns with other work suggesting that reducing the activity of CRF expressing neurons reduces alcohol-related behaviors. Other work targeting CRF neurons using a chronic exposure model showed increased excitability of a putative non-projecting population (those that do not project to the VTA or lateral hypothalamus; Pati et al., 2020), highlighting specific subpopulations (CRF only) and circuits (local versus projecting) that may be involved in these behaviors. The literature suggests that an alternate, distinct subpopulation of BNST neurons may play an opposing role in alcohol consumption (Kash & Winder, 2006). Neuropeptide Y (NPY) signaling within the BNST reduces alcohol consumption (Pleil et al., 2015). This is of particular interest because SST neurons can co-localize with NPY neurons (Epelbaum et al., 1994) – although elsewhere, they co-localize with Dynorphin expressing neurons (Kim et al., 2017) – highlighting for a need to comprehensively characterize SST neurons in the BNST. Overlap between SST and NPY, however, is congruent with the overall hypothesis that activation of both/either of these subpopulations reduces alcohol consumption. Indeed, our work provides preliminary evidence that manipulation of the SST subpopulation in the dorsolateral BNST can augment drinking behavior in the drinking in the dark model. This complements changes in SST excitability we found in our previous work utilizing intermittent access to and forced withdrawal from alcohol (Dao et al., 2020). Though this work suggests BNST SST neurons are a promising target for AUD, much work needs to be done to characterize the role of these neurons in different models of alcohol, points along the trajectory to development of AUD, and the role differing SST circuits may play. Emphasis needs to be placed on characterizing these SST neurons, in terms of their co-expression and overlap with known alcohol-influencing peptidergic populations, as well as the sources of their activation and their projection targets.

### 4.3 The BNST, SST, and anxiety

Importantly, both the BNST and SST broadly play a larger role than just in the modulation of alcohol consumption. The BNST has rich connectivity with other important structures in the limbic system and plays a role in behaviors such as sustained fear, generalized anxiety disorder, post-traumatic stress disorder, feeding, and social anxiety (Flanigan & Kash, 2020; Hardaway et al., 2015; Sullivan et al., 2004). Upregulation of SST neurons throughout the brain has been shown to ameliorate anxiety-like and depressive-like behavior (Fuchs et al., 2017). Indeed, SST neurons have been reasonably well characterized in cortical circuits (Urban-Ciecko & Barth, 2016; Cummings and Clem, 2020; Dao et al., 2021)_where they are found to have unique electrophysiological properties and can modulate a variety of behaviors. SST neurons have been less thoroughly investigated in subcortical regions, however. Our work here did not show any clear role of BNST SST neurons in anxiety-like behaviors – however, our investigations were limited to chemogenetic activation of dorsolateral SST neurons, with a limited number of behavioral manipulations. Our injections did not target ventral BNST neurons which may play a unique role in these behavioral phenotypes.

Previous work has demonstrated that Gq-activation of GABAergic neurons in the BNST confers an anxiogenic-like profile (Mazzone et al., 2018). The comprehensive assessment in Mazzone et al. captured a much wider population of neurons – notably, both dorsal and ventral BNST (whereas our current work more greatly profiles dorsal BNST), and all GABAergic neurons (whereas our work captures the SST-expressing subset of this population). It is possible that modulation of anxiety-like behavior requires the synchrony of many GABAergic neurons, of which the SST population may be a part. Recent work has shown that a largely SST-expressing BNST population that projects to the nucleus accumbens can modulate anxiety (Xiao et al., 2020). However, this work used more intensive optogenetic activation strategies, which may not be captured with DREADD strategies. Alternatively, it may be that manipulation of pathway-specific SST neurons in the BNST is necessary to modulate anxiety, whereas overall activation of these neurons contributes to opposing pathways and otherwise dilutes behavioral effects. In addition, other work suggests that BNST peptide-expressing populations may be modulated by insults such as stress in a strain-dependent manner (Pleil et al., 2012). A greater characterization of SST pathways, and overlap with other peptidergic populations in the BNST, may provide better clarity. Further investigations should dive deeper into the role of BNST SST neurons in anxiety (if any), including following dependent models of alcohol consumption such as intermittent access (Holleran et al., 2016; Dao et al., 2020).

### 4.4 SST neurons in the BNST - projections and overlapping populations

Little is known about the projection patterns and expression of BNST SST neurons. SST neurons are co-expressed with NPY neurons in the BNST of rodents (Kash et al., 2015, Epelbaum et al., 1994) and this aligns with the overall ‘anti-drinking’ role we describe here. New work by Bruzsik et al. (2021) found that BNST SST and CRF neurons target distinct postsynaptic targets, and the authors suggest they likely have little overlap (Bruzsik et al., 2021). Importantly, this is consistent with our hypothesized circuit here. This circuit is activated (at least in part) by SST expressing neurons within the CeA (Ahrens et al., 2018). In addition, recent publications suggest that posterior BNST SST neurons may project to regions of the hypothalamus (Barbier et al., 2021). Importantly, this paper explored SST neurons in chronic stress, suggesting subpopulations of BNST SST neurons may be more relevant for anxiety-related behaviors than those we captured with a broad DREADD approach. New work has suggested that SST neurons in the BNST do not only serve as ‘local’ GABAergic inhibition and project to the nucleus accumbens (Xiao et al., 2020) and a careful accounting of BNST SST neuronal morphology and circuit connections is needed.

### 4.5 Sex Differences

Our work found sex differences in the ability to shift binge alcohol consumption using chemogenetic approaches. This is consistent with a significant literature indicating sexual dimorphism found in the BNST, and importantly, that this region can be influenced by gonadal hormones (Allen & Gorski, 1990). Up until recently, much of the substance use literature has focused exclusively on male mice, and the literature has increasingly highlighted the need to explore both sexes in AUD and substance use research more broadly (Jury et al., 2017). Withdrawal from other drugs of abuse (morphine) differentially regulates anxiety-like behavior (Bravo et al., 2020) and GABAergic transmission in the BNST (Luster et al., 2020). In addition, other neurotransmitter systems appear to differentially modulate BNST driven behaviors such as anxiety (Marcinkiewcz et al., 2019). Coupled with the known differences in males and females in alcohol consumption and AUD development (Grant et al., 2015), there is a need to comprehensively characterize BNST SST neuronal differences in males and females both at a basal state and following models of disease state perturbations. Our work provides preliminary data surrounding the SST neuronal system in the BNST in control mice, suggesting a greater need for research here. We demonstrate a difference in overall number of SST neurons in both the dorsal and ventral BNST of males versus females, though this can be accounted for by the relatively larger amygdala size of male mice. Future work should investigate whether this population of neurons sends differing projections in the two sexes. Though our intention here was to include both sexes and not explore cycling sex hormones as an independent variable, future work should do so.

## 6. CONCLUSIONS

Here, we demonstrate that chemogenetic activation of BNST SST neurons reduces binge drinking in female mice with no major effects on anxiety-like behavior. This provides preliminary evidence for these neurons as a promising target for the treatment of AUD. Future work should further tease apart SST activation and connectivity in this region, as well as the effects of SST peptide signaling itself.

## AUTHOR STATEMENT

MSN and NCD contributed equally to this work. MSN: conceptualization, data curation, formal analysis, wrote manuscript, edited manuscript. NCD: data curation, formal analysis, wrote manuscript, edited manuscript. DLM: data curation, formal analysis. KRG: data curation, formal analysis. WDS: data curation. JBM: data curation. AS: data curation, formal analysis. DFB: data curation. KDM: data curation. NAC: conceptualization, data curation, formal analysis, wrote manuscript, edited manuscript. The authors declare no competing interests.

## ACKNOWLEDGEMENTS

This work was supported by The National Institutes on Alcohol Abuse and Alcoholism of the National Institutes Health (1R21AA028088 to NAC; P50AA017823 to NAC), The Brain and Behavior Research Foundation (NARSAD Young Investigator Award to NAC), and The National Institute of General Medical Sciences (T32GM108563 to ARS)

